# Automatic Localization of the Subthalamic Nucleus on Patient-Specific Clinical MRI by Incorporating 7T MRI and Machine Learning: Application in Deep Brain Stimulation

**DOI:** 10.1101/322230

**Authors:** Jinyoung Kim, Yuval Duchin, Reuben R. Shamir, Remi Patriat, Jerrold Vitek, Noam Harel, Guillermo Sapiro

## Abstract

Deep Brain Stimulation (DBS) of the subthalamic nucleus (STN) has shown clinical potential for relieving the motor symptoms of advanced Parkinson’s disease. While accurate localization of the STN is critical for consistent across-patients effective DBS, clear visualization of the STN under standard clinical MR protocols is still challenging. Therefore, intraoperative microelectrode recordings (MER) are incorporated to accurately localize the STN. However, MER require significant neurosurgical expertise and lengthen the surgery time. Recent advances in 7T MR technology facilitate the ability to clearly visualize the STN. The vast majority of centers, however, still do not have 7T MRI systems, and fewer have the ability to collect and analyze the data. This work introduces an automatic STN localization framework based on standard clinical MRIs without additional cost in the current DBS planning protocol. Our approach benefits from a large database of 7T MRI and its clinical MRI pairs. We first model in the 7T database, using efficient machine learning algorithms, the spatial and geometric dependency between the STN and its adjacent structures (predictors). Given a standard clinical MRI, our method automatically computes the predictors and uses the learned information to predict the patient-specific STN. We validate our proposed method on clinical T_2_W MRI of 80 subjects, comparing with experts-segmented STNs from the corresponding 7T MRI pairs. The experimental results show that our framework provides more accurate and robust patient-specific STN localization than using state-of-the-art atlases. We also demonstrate the clinical feasibility of the proposed technique assessing the post-operative electrode active contact locations.

## 1. Introduction

Deep brain stimulation (DBS) is a neuromodulation intervention for relieving the motor symptoms of Parkinson’s disease (PD), dystonia, and Essential tremor, among others [Dormont et al., 2010; Kim et al., 2010; Krack et al., 2010; Limousin et al., 1998; Mallet et al., 2007; Patel et al., 2008; The Deep-Brain Stimulation for Parkinson’s Disease Study Group, 2001; Volkmann, 2007]. In particular, DBS of the subthalamic nucleus (STN) has been shown to be an effective symptom’s treatment for advanced PD [Dormont et al., 2010; Limousin et al., 1998; The Deep-Brain Stimulation for Parkinson’s Disease Study Group, 2001; Volkmann, 2007].

Accurate 3D positioning of the chronic electrode within the STN is critical for the success of the DBS surgery, as its efficacy and adverse effects are highly correlated with the electrode’s location [Hamid et al., 2005; Kerl et al., 2012; Mallet et al., 2007; Patel et al., 2008; Starr et al., 2002]. Thus, precise identification of the STN (lens-shaped) of individual patients facilitates the DBS planning (and post-op programming). However, such identification in clinical settings still remains challenging due to its small size (approximately 6×4×5mm) and the ambiguous border with neighboring regions [Abosch et al., 2010].

Both direct- and indirect-targeting is often incorporated to estimate the 3D STN location and shape. Direct identification of the STN and its surrounded regions (e.g., substantia nigra (SN) and zona incerta) is possible using various MRI modalities where these structures’ appearance is hypo-intense such as T_2_W, fluid attenuated inversion recovery (FLAIR), fast gray matter acquisition T_1_ inversion recovery (FGATIR), and susceptibility weighted imaging (SWI). However, the distinction amongst the neighborhood structures is often unclear on the clinical MRI, and the STN appears in only one or two slices, thereby resulting in sub-optimal targeting within the 3D STN (positive effects), and relative to adjacent structures of the STN (potential negative/side effects). There are efforts underway to directly visualize the STN via MRI reconstruction methods such as susceptibility weighted phase imaging or quantitative susceptibility mapping (QSM) [Chandran et al., 2015; Liu et al., 2013; Rasouli et al., 2017]. Moreover, recent studies have proposed automatic methods to segment the STN on these contrast enhanced MR sequences [Garzón et al., 2017; Milletari et al., 2017; Visser et al., 2016b]. While early work on QSM shows promising results, uncertainty in susceptibility estimates under different acquisition protocols needs to be further investigated [Lauzon et al., 2016] and the overall performance needs to be validated on large-scale clinical data.

The indirect STN localization approach refers to the selection of targets based on nearby anatomical landmarks and then the computation of consensus coordinates estimates in relation to these landmarks. The most common method is to select the anterior-and posterior-commissure anatomical landmarks on the clinical MRI, define the midline and the mid commissural point, and use the consensus coordinates ±12mm lateral, 4mm posterior, and 5mm inferior to the mid commissural point [Starr et al., 2002]. This is used as an initial estimation for the location of the STN that is later refined based on the neurosurgeon’s expertise and preferences. However, this consensus method does not account for the obvious variability in the patients’ anatomy [Daniluk et al., 2010; Kerl et al., 2012]. Moreover, it was reported that there is significant inter-surgeon variability in the selection of anterior-and posterior-commissure points, which has a substantial effect on the localization of targets using this indirect method [Pallavaram et al., 2008]. For these reasons, indirect targeting is almost always complemented by other refinement methods.

Atlas-based approaches have been proposed to improve targeting accuracy. Some studies show promising results for the visualization of the STN for DBS surgery on 3 Tesla (T) MR datasets. Patch based label fusion methods were used for segmenting the STN and its adjacent structures using multimodal 3T MRIs [Haegelen et al., 2013; Xiao et al., 2014a]. Xiao et al. [2014b] analyzed the morphometric variability of the STN obtained by a majority voting label that was augmented on 3T MRIs of advanced PD patients. D’Albis et al. [2015] provided a pipeline for DBS planning and post-operative validation and adopted an atlas-based segmentation in the surgical planning flow. Post-operative active contacts’ clusters projected onto the CranialVault atlas were used for DBS target prediction in [Pallavaram et al., 2015]. A probabilistic approach to map DBS electrode coordinates onto the MNI space was presented in [Horn et al., 2017a]. More recently, to achieve anatomical precision in the MNI space, a histological atlas, described in Chakravarty et al. [2006], was merged with subcortical atlases based on multiple contrast 3T MR sequences from PD patients [Xiao et al., 2017] and high resolution multimodal MRIs and structural connectivity data [Ewert et al., 2017].

While encouraging initial results were obtained with these techniques, the obtained targeting accuracy is oftentimes insufficient, due in part to the large per-patient STN variability, to fully ensure DBS treatment efficacy and safety. Based on patient-specific clinical data, further revisions of the approximated lead location are often required when using these atlas-based approaches.

Microelectrode recordings (MER) are often incorporated to define the precise location of the STN for correct placement of the electrode within the targeted structure. These electrophysiological measurements require significant team expertise and extend the surgery time. MER are often complemented with patient’s behavioral feedback while the subject is awake [Abosch et al., 2010]. Note that brain shift has been shown to increase with the length of the procedure (e.g., resulting from extended MER), causing further challenges. Others have suggested that while the incidence of brain shift is infrequent, it is also unpredictable [Halpern et al., 2008; Ivan et al., 2014; Petersen et al., 2010]. Lack of standardization and variability in how individual centers use MER emphasizes the importance of developing additional standard and objective approaches for targeting the STN.

With recent advances in ultrahigh magnetic fields hardware and acquisition protocols, 7T MR imaging techniques now allow the direct identification of small and complex anatomical structures, including the 3D STN, thanks to its superior contrast and high resolution [Abosch et al., 2010; Cho et al., 2011; Kerl et al., 2012]. Furthermore, 7T MRI has already facilitated the study of connectivity within the basal ganglia and thalamus and enabled the subdivision of the STN into motor, associative, and limbic sub-regions [Abosch et al., 2010; Lenglet et al., 2012; Plantinga et al., 2016]. Keuken et al. [2013] investigated structural change of the STN using atlases based on 7T MRI in different age groups (healthy subjects). Probabilistic atlas maps obtained from multiple 7T MR contrasts were used for analysis of anatomical variability on subcortical structures [Keuken et al., 2014]. Wang et al. [2016] generated the ultrahigh-field atlas of the STN using the 7T T_1_-weighted (T_1_W) and T_2_-weighted (T_2_W) MRI. For use with clinical data, Plassard et al. [2017] created atlases based on the 7T MRI of healthy subjects and used them to segment subcortical structures with 3T contrast enhanced MR sequences of the same subjects. Moreover, a high quality 7T atlas obtained from elderly subjects was registered onto the 3T MRI template averaged on PD patient’s data [Milchenko et al., 2018]. Automated methods to segment brain subcortical structures using 7T MRI have been proposed, leveraging sufficient intensity information from multiple MR contrasts [Kim et al., 2014; Visser et al., 2016a; Visser et al., 2016b]. The clinical feasibility of 7T MRI for localization of the STN has been previously demonstrated [Duchin et al., 2012]. More recently, the U.S. Food and Drug Administration (FDA) cleared the first 7T MRI system (The Magnetom Terra, Siemens Medical Solutions) for clinical use. However, 7T MR machines are still rare in current clinical practice and are associated with significant infrastructure costs [Plantinga et al., 2014]. Therefore, the STN still needs to be localized on the ubiquitous clinical platforms of 1.5T or 3T MRI.

In this work, we propose to incorporate a high-quality 7T MR dataset for training machine learning methods to statistically model geometrical dependencies of the subcortical structures (the framework proposed here is named “7T-ML”). We demonstrate that this approach facilitates the accurate prediction of the patient-specific STN location and shape on standard clinical MRI, [^1^ The framework and algorithm here described are components of the patented and FDA cleared patientspecific STN visualization tool developed by Surgical Information Sciences, Inc. [Harel and Sapiro, 2016; Sapiro et al., 2017]. A preliminary work was presented at conferences [Kim et al., 2015a; Kim et al., 2015b; Kim et al., 2015c]. The scope of this study is the validation and analysis of the method on a large scale clinical data, and comparison of the proposed method with available state-of-the-art methods.] thereby enjoying the advantages of high-quality data without the need for a 7T machine, or any additional hardware. The proposed 7T-ML framework is extensively compared with ground truth obtained from 7T MRI of the same subjects and with state-of-the-art atlas-based results (comprising histology, healthy population, Parkinson’s population, and elderly, thereby introducing high variability in the testing and validation), showing the significantly improved performance, which is critical for accurate targeting, DBS lead localization, and clinical outcomes. The algorithm’s relevance for DBS in particular and neuromodulation in general is demonstrated with the accurate visualization of active contacts from real DBS surgeries.

## 2. Methods

### 2.1 Overview

We create an annotated dataset of multiple pairs (same subject) of clinical (1.5T or 3T) and 7T MRIs along with the segmented (labeled) subcortical structures of interest. The 3D STN was segmented on the 7T MRI, but not on the clinical image (where as discussed above, is not clearly visible). Subcortical structures in the vicinity of the STN were also segmented on the 7T MRI (hereafter named “predictors”), these are visible on the clinical image as well. This dataset was used for learning the geometric relationships between the STN and its predictors. The dataset containing the clinical and 7T MRIs along with the segmented structures is referred hereafter as the “training set.” Given a new patient’s clinical MRI (no 7T images for this patient), the algorithm automatically detects the predictors and then computes the patient-specific, i.e., patient’s own, 3D STN location and shape. This is based on the learned information from the training set. To evaluate and validate the quality of the automatic STN localization, the results are compared with the STN segmented by experts on the 7T MRI (the standard data split between training and testing set is used here), and further clinically validated with active contact positions obtained from the post-operative CT of the same patient. Comparisons with existing literature are provided as well, showing a significant improvement on accuracy and robustness, both critical for patient-specific STN DBS and neuromodulation targeting.

### 2.2 Database and Preprocessing

7T MRI and its corresponding (same subject) clinical MRI (1.5T or 3T) of 80 subjects were used in this study under approval of the Institutional Review Board at the University of Minnesota. Demographic and clinical details for the subjects are presented in Table 1. Table 2 presents the MRI modality and resolution of the 7T and clinical data that were used in this work.

**Table 1:** Demographic and clinical details of the used 80 subjects.

| Gender | | Diagnosis | | | Age<br>(mean ± standard deviation) |
| --- | --- | --- | --- | --- | --- |
| Male | Female | Essential Tremor | Parkinson's Disease | Normal Control |  |
| 61 | 19 | 12 | 56 | 12 | 60±13.8 |

**Table 2:** MRI modality and resolution of 7T and clinical data used in our 7T-ML.

| Magnetic field strength | 7T |  | 1.5T | 3T |
| --- | --- | --- | --- | --- |
| MRI modality | T<sub>2</sub>W | SWI | T<sub>2</sub>W | T<sub>2</sub>W |
| Resolution | 0.39x0.39x1.0mm<br>0.39x0.39x2.0mm | 0.39x0.39x0.8mm | 0.5x0.5x2.0mm<br>0.72x0.72x2.0mm | 0.55x0.55x2.0mm |

Experts in the team manually segmented the STN and its predictors on the 80 pairs of 7T T_2_W and SWI MRIs, and carefully cross-validated segmentation results. The 7T data acquisition protocol, preprocessing, and manual segmentation are detailed in Lenglet et al. [2012] and omitted here for brevity. For illustration purposes, manual delineation of the STN and SN on a pair of 7T T_2_W and SWI MRIs is visualized in Fig. 1. Duchin et al. [2018] also demonstrated that the 7T MRI based STN segmentation highly agrees with MER data (see also Shamir et al., [2018]). Hypo-intense regions that are spatially adjacent with the STN are considered as predictors. Such subcortical structures are well visualized both on the 7T and clinical MRIs [Cho et al., 2011].

**Figure 1:**
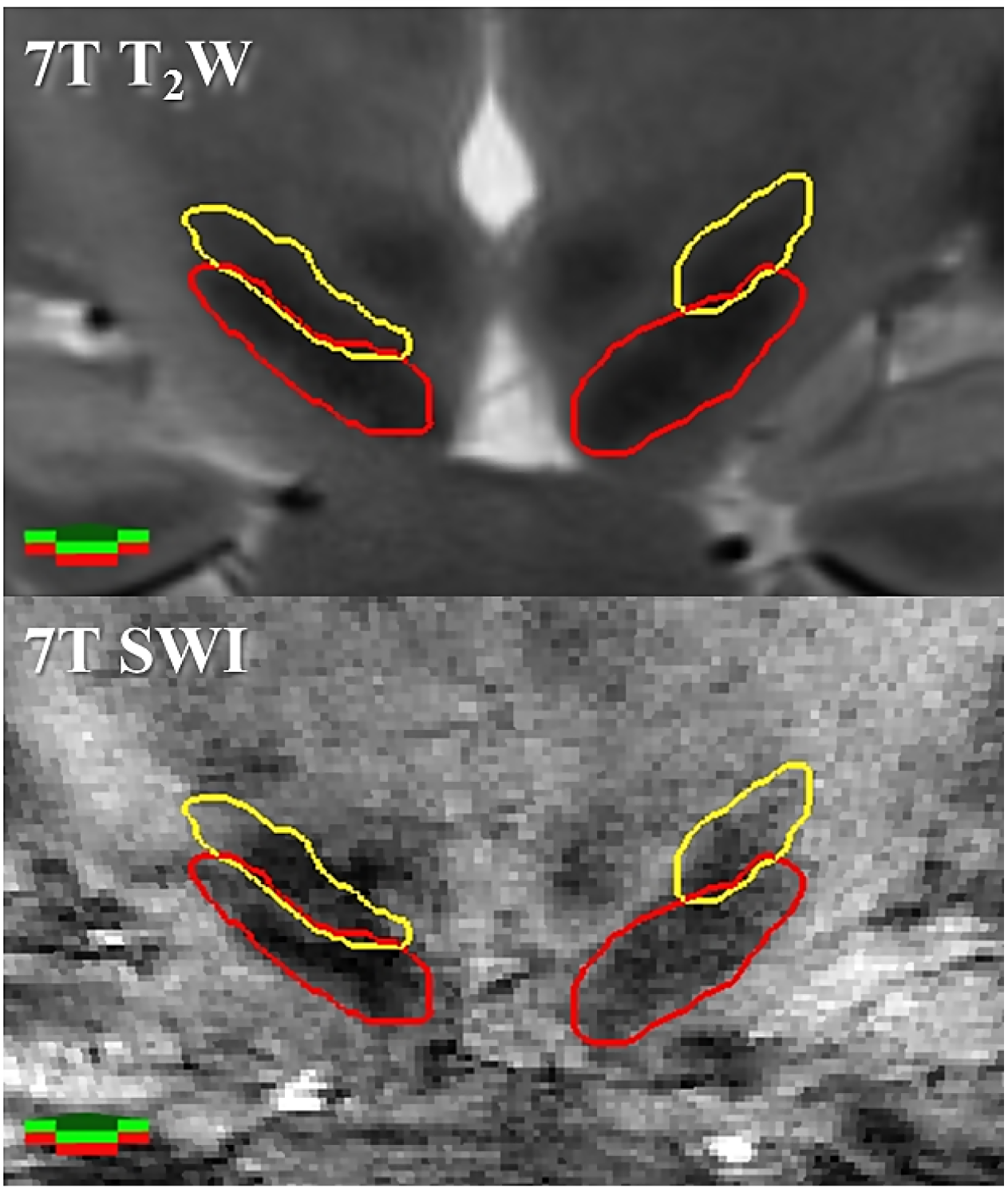
Direct visualization of SN and STN on a coronal 7T T_2_W image (top) registered onto SWI, coronal 7T SWI (bottom). The red and yellow represent contours of manually segmented SN and STN, respectively. Green arrows are toward the anterior direction.

The 7T T_2_W MRIs were co-registered to their clinical T_2_W MRI counterparts using the FSL FMRIB’s Linear Image Registration Tool [Jenkinson and Smith, 2001], and the structures segmented on the 7T images were transformed accordingly (hereafter “7T priors”). Data acquisition, pre-processing and co-registration for 7T and clinical MRI (1.5T or 3T) used in this study were performed following protocols described in Duchin et al. [2012].

The MRI pairs and the 7T priors were all stored in a database for retrieval upon the introduction of a new patient’s clinical MRI on which the STN needs to be localized. 46 datasets of 7T and 1.5T MRIs pairs of patient groups were used for training and 34 datasets of 7T and corresponding 1.5T or 3T MRIs of patients and normal control subjects were added in our database later for further validation. A subset of 15 PD patients in the training set were selected to further study in detail various factors that affect the performance of our algorithm. Table 3 presents clinical details on to magnetic field strength of the clinical images in the training and validation sets. Note that 3T MRIs were not used at all to train the algorithm, only for validation.

**Table 3:** Clinical details and magnetic field strength of clinical images in the training and validation sets (80 subjects).

| Dataset | Training set | Validation set |  |
| --- | --- | --- | --- |
| Magnetic field strength | 1.5T | 1.5T | 3T |
| The number of patients | 46 (15) | 10 | 24 |
| Essential Tremor | 10 | - | 2 |
| Parkinson's Disease | 36 (15) | 10 | 10 |
| Normal Control | - | - | 12 |

( ) indicates the number of PD patients for in-depth study

Given a new clinical T_2_W MRI, we first compute its affine registration with all 46 clinical T_2_W MRIs in the training set within our database. Intensity similarity scores are measured by calculating the mutual information between mid-brain regions of the clinical query image and the registered images in the training set [Wells III et al., 1996]. About 30% of training images sorted by the similarity scores are empirically selected as the most similar sets to reduce biases [Aljabar et al., 2009], and then they are nonlinearly registered onto the clinical query image. The Advanced Normalization Tools [Avants et al., 2011a; Avants et al., 2011b; Avants and Gee, 2004] was used for global affine registration (with the mutual information cost metric) and local nonlinear registration of the basal ganglia region (with crosscorrelation cost metric, SyN option, and bspline interpolation). Then, we resample the clinical query image to a 0.5mm isotropic resolution and transform the selected datasets into the query image coordinates system. As a result, the clinical query data and the 7T priors in the training set are now in a common coordinate system.

### 2.3 7T-Machine Learning based STN Prediction

The geometric relationship between the 3D STN and its predictors on the training set was analyzed using a regression-based shape prediction approach [Baka et al., 2011; Blanc et al., 2012; Rao et al., 2008].

We first automatically segment the predictors on a query clinical T_2_W MRI, which are observable subcortical structures near the STN, later used to predict it. To this end, the training dataset is incorporated in a unified framework of active shape model and active appearance model algorithms [Cerrolaza et al., 2012; Cerrolaza et al., 2015; Cootes et al., 1995; Frangi et al., 2002; Heimann and Meinzer, 2009; Matthews and Baker, 2004; Sung et al., 2007; Tzimiropoulos and Pantic, 2013]. Then, we apply the regression-based shape prediction, incorporating the computed predictors on the query clinical image and the geometric relationship learned from the STN and the segmented predictors on the training dataset.

More specifically, 3D shapes of the predictors and the STN are represented as the coordinates of surface points (vertices in a mesh), in correspondence, across random subsets of the most similar training sets previously selected (registered onto the clinical query image). The poses of the structures are extracted using the generalized Procrustes analysis [Cootes et al., 1992; Gower, 1975], and their shape variations are then modeled using kernel principal component analysis [Guo, 2010; Rathi, 2006]. A bagging procedure [Breiman, 1996] is applied in the partial least squares regression technique [Abdi, 2010; Krishnan et al., 2011; Wold, 1982; Wold et al., 2001] to learn the dependency between the STN and its corresponding predictors (in the pose and shape feature space) from these randomly selected subsets of the most similar training sets [Kim et al., 2015b]. We also investigated a regression forest model [Breiman, 2001; Criminisi et al., 2013] for finding the non-linear mapping between the STN and its predictors [Kim et al., 2015c]. Given the computed (visible) predictors in a new patient clinical data, the 3D STN binary volume is predicted by exploiting this learned spatial relationships. See Kim et al. [2015b] and Kim et al. [2015c] for additional technical details.

Such an ensemble approach with equal weights does not consider the influence of each training subset on the final confidence map of the predicted STN. If some training subsets are more influential than others with respect to the specific patient’s STN prediction, the prediction accuracy can be further improved by increasing their weights/relevance. To estimate the contribution of each training subset to the prediction, we investigate non-linear relationships between pose-related features [Kim et al., 2015a] and the prediction accuracy from various subjects and corresponding training subsets. This is done using a random forest model [Breiman, 2001]. The global error score, namely a weighted sum of all the geometric measures, was used to determine the influence-weight of each training subset on the final prediction [Kim et al., 2015a]. Given new features from a query patient and each training subset, the error scores are estimated using the learned information. An ensemble of predictions from the random subsets with larger weighting (i.e., more influential sub-atlases for the prediction) yields lower error scores. The weighted ensemble also provides a final confidence map on the query patient. We refer to this procedure as “robust prediction” to differentiate it from “ensemble,” that simply incorporates equal weights for all the subsets. See additional details and explicit parameters’ values in Kim et al. [2015a].

### 2.4 Validation

To evaluate the performance of the proposed 7T-ML method, we used clinical data (1.5T or 3T MRI) from 80 subjects (46 cases using a leave-one-out approach on the training set and additional 34 naïve cases for validation, not present at all in the training). One clinical MRI was selected as a query image at each iteration. When the query image was from the training set, all its associated data (segmented structures and other MRIs) was excluded from the 46 training sets. The STN was then automatically computed on the query image using the proposed framework. This was done with automatically segmented (on the clinical data) predictor structures, bagged partial least squares regression, and uniform weights of 100 training subsets. The current implementation utilizes Amazon Web Services to facilitate the large scale parallel processing.

Our 7T-ML method is compared with the following publicly available STN atlases in the literature to validate its reliability for patient-specific clinical MRI based STN-DBS targeting: 1) 7T MRI STN atlas for use with 3T MRI (7T3T) [Milchenko et al., 2018]; 2) ultrahigh-field atlas (UHFA) [Wang et al., 2016]; 3) DBS intrinsic template atlas (DISTAL) [Ewert et al., 2017; Horn et al., 2017b]; and 4) population-averaged atlas that was made with 3T MRI of 25 PD patients (MNIPD25) [Xiao et al., 2017]. Data acquisition and template images for these atlases are summarized in Table 4. This is a very comprehensive comparison since these atlases include 7T data, histology, healthy subjects, PD patients, and elderly. To this end, we computed the transformation between the atlas MRI templates and the clinical T_2_W MRIs across 80 subjects, and standard STN atlases were accordingly transformed to the clinical data. For a fair comparison, we applied the same approach as the inter-patient registration between database clinical images and a clinical query image in our proposed framework. It is a multi-step registration (global affine registration and local nonlinear registration on the basal ganglia region) that was adjusted based on a long trial and error process.

**Table 4:** Data acquisition and template images for tested state-of-the-art atlases.

| Atlases | 7T3T<br>(Milchenko et al., 2018) | UHFA<br>(Wang et al., 2016) | DISTAL<br>(Ewert et al., 2017) | MNIPD25<br>(Xiao et al., 2017) |
| --- | --- | --- | --- | --- |
| Data on which<br>the STN atlas<br>was obtained | 7T T<sub>2</sub>W MRI<br>(elderly normal) | 7T T<sub>2</sub>W MRI<br>(normal) | 3T T<sub>1</sub>W/T<sub>2</sub>W/proton<br>density MRI (normal),<br>histology, and<br>DWI<br>(normal/PD patients) | 3T T<sub>2</sub>*W MRI<br>(PD patients) |
| Provided STN<br>atlas template<br>image | 3T T<sub>2</sub>W MRI in ICBM<br>MNI152 space<br>(PD patients) | 7T T<sub>2</sub>W MRI<br>(normal) | 3T T<sub>2</sub>W MRI in<br>ICBM 2009b Asym<br>MNI152 space<br>(normal) | 3T T<sub>2</sub>*W MRI in<br>ICBM MNI152<br>space (PD patients) |

Additionally, we provide the STN segmentation results obtained with a conventional approach based on intensity information on the image - the active shape model and active appearance model framework (also used for predictor segmentation in our 7T-ML) [Sung et al., 2007; Tzimiropoulos and Pantic, 2013]. This also motivates a novel approach in challenging situations, e.g., when the STN borders are not visible.

The STN manually segmented on the 7T MRI and transformed onto the clinical MRI pair of the same subject was used as “ground truth.” The following measures were first calculated to compare our 7T-ML and standard atlases with the 7T manual ground truth STN: 1) distance between the centers of mass; 2) mean Euclidean distance of surface points in correspondence; 3) Dice coefficient (DC), which is a normalized overlap measure between two co-aligned binary datasets, and; 4) volume of the STN (added here for completeness). As suggested in Shamir et al. [2009], 2mm accuracy is considered an acceptable threshold for neurosurgical and neuromodulation applications. Therefore, we counted the number of cases that are less than 2mm centers of mass distance as a clinically relevant prediction accuracy measure. Our 7T-ML STN confidence map and standard atlases-based STN were normalized to [0, 1] and binarized with threshold values between 0.3 and 0.4 to eliminate resampling artifacts, thereby avoiding a bias in the measures induced by unexpectedly large volumes. A one-way analysis of variance (ANOVA) was calculated for each measure and a multiple comparison correction was performed to estimate the methods. Then, post hoc tests with Tukey’s honest significant difference were executed.

To demonstrate the relevance of the high accuracy of the proposed 7T-ML approach for DBS and neuromodulation, we also computed the distance between the DBS chronic electrode’s active contacts and the predicted STN’s centers of mass. The center of mass is used here for reference, providing intrinsic distances for proper and consistent comparison (also across subjects); it should be noted that the STN’s center of mass was not considered as the optimal target point. Pallavaram et al. [2015] measured the distance between active contacts and the estimated target position (using different targeting protocols, e.g., stereotactic coordinates corresponding to the center of the STN’s motor territory). For this, pre-operative clinical MRI and the STNs computed from each method are co-registered onto the post-operative CT of the same patient (where the implanted electrode is detected). We extensively examined the registration results to ensure the correct transformation of the STN.

Furthermore, to compare across different approaches the active contacts location of individual patients, we used a best-fit ellipsoid representation of the individual’s STN. Then, we divided the space into eight regions (in posterior/anterior, medial/lateral, and superior/inferior axes) and counted the number of active contacts in each region (for all patients for which we have available data). We defined an active contact as “in” a specific region of the STN if its center of mass was in that region. Note that in contrast to common averaging techniques where very different shapes are (often significantly) deformed into a common coordinate system, this is a patient-specific intrinsic measurement, each individual STN is split into its own 8 regions, and then votes are accumulated. This study demonstrates the sub-region accuracy of the proposed visualization/targeting approach that is critical both to localize the active DBS contacts and for future studies on the clinical targeting value of different STN sub-regions. The clinical data used for the post-operative analysis in the training and validation sets is summarized in Table 5. We used 38 electrode’s lead images (with information available on 31 active contacts, the others had missing clinical data) reconstructed from the post-operative CT of 30 PD patients out of the total 80 subjects. These represent all the subjects from which we could retrieve from the available clinical records both postoperative data and outcome information. This study was repeated also for the location of non-active contacts for completeness.

**Table 5:** Summary of patient data used for the post-operative analysis in the training and validation sets.

| Dataset | Training set | Validation set |
| --- | --- | --- |
| The number of patients | 46 | 34 |
| The number of patients for post-operative analysis | 15 | 15 |
| Unilateral STN-DBS | 12 (11/12) | 10 (9/10) |
| Bilateral STN-DBS | 3 (5/6) | 5 (6/10) |

( ) represents the number of available active contacts out of total electrode reconstruction image

Lastly, we selected from the training set 15 PD patients whose clinical 1.5T T_2_W MRIs contain a whole head image in order to investigate the different factors that affect the STN prediction in our proposed 7T-ML framework. For this in-depth analysis, we computed the STN under various setups: 1) *Predictors*: manual or automatic segmentation; 2) *Regression methods*: bagged partial least squares regression or random forest; 3) *Ensemble sizes*: 10, 100, and 200; and 4) *Weighting methods*: uniform or non-uniform weights on training subsets based on their contribution to the prediction accuracy.

## 3. Results

Our proposed 7T-ML and standard atlases-based STNs are compared with the 7T manual ground truth STN on 80 (160 STNs) subjects’ clinical data, Table 6 and Fig. 2. The 7T-ML STNs were significantly closer to the 7T manual ground truth STNs in comparison to the atlas-based results (p<0.0001; ANOVA). Our results demonstrate the high accuracy and consistency of the proposed 7T-ML. Compare the 1.25±0.60mm, 2.37±1.74mm, 2.94±1.49mm, 3.50±3.57mm, and 4.25±3.33 average centers of mass distance between the STN that was computed from the 7T-ML, 7T3T, UHFA, DISTAL, and MNIPD25 atlases, respectively, and the ground truth STN (p<0.0001; ANOVA). Note that the average error in the 7T-ML STN is close to half of the clinical data slice thickness (2mm), which is roughly a lower bound to the possible segmentation accuracy. Furthermore, 89.4% (143/160), 53.8% (86/160), 28.1% (45/160), 51.9% (83/160), and 24.4% (39/160) of the STNs computed from the 7T-ML, 7T3T, UHFA, DISTAL, and MNIPD25 atlases, respectively, were less than 2mm from the ground truth center of mass (acceptable maximal error).

**Table 6:**
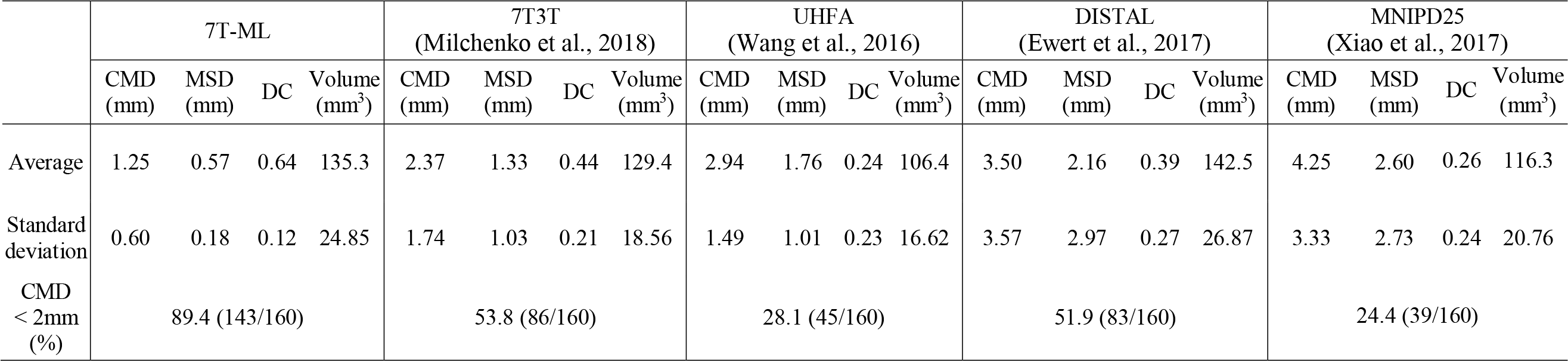
Quantitative comparison of the STNs obtained using the proposed 7T-ML and the 7T3T, UHFA, DISTAL, MNIPD25 atlases. The various methods were compared to the 7T manual ground truth STNs of 80 patients (160 STNs). A one-way ANOVA and post hoc test (with Tukey’s method) showed that the proposed 7T-ML is significantly better than atlas-based methods in each measure (p<0.0001). CMD: centers of mass distance. MSD: mean distance of surface points.

**Figure 2:**
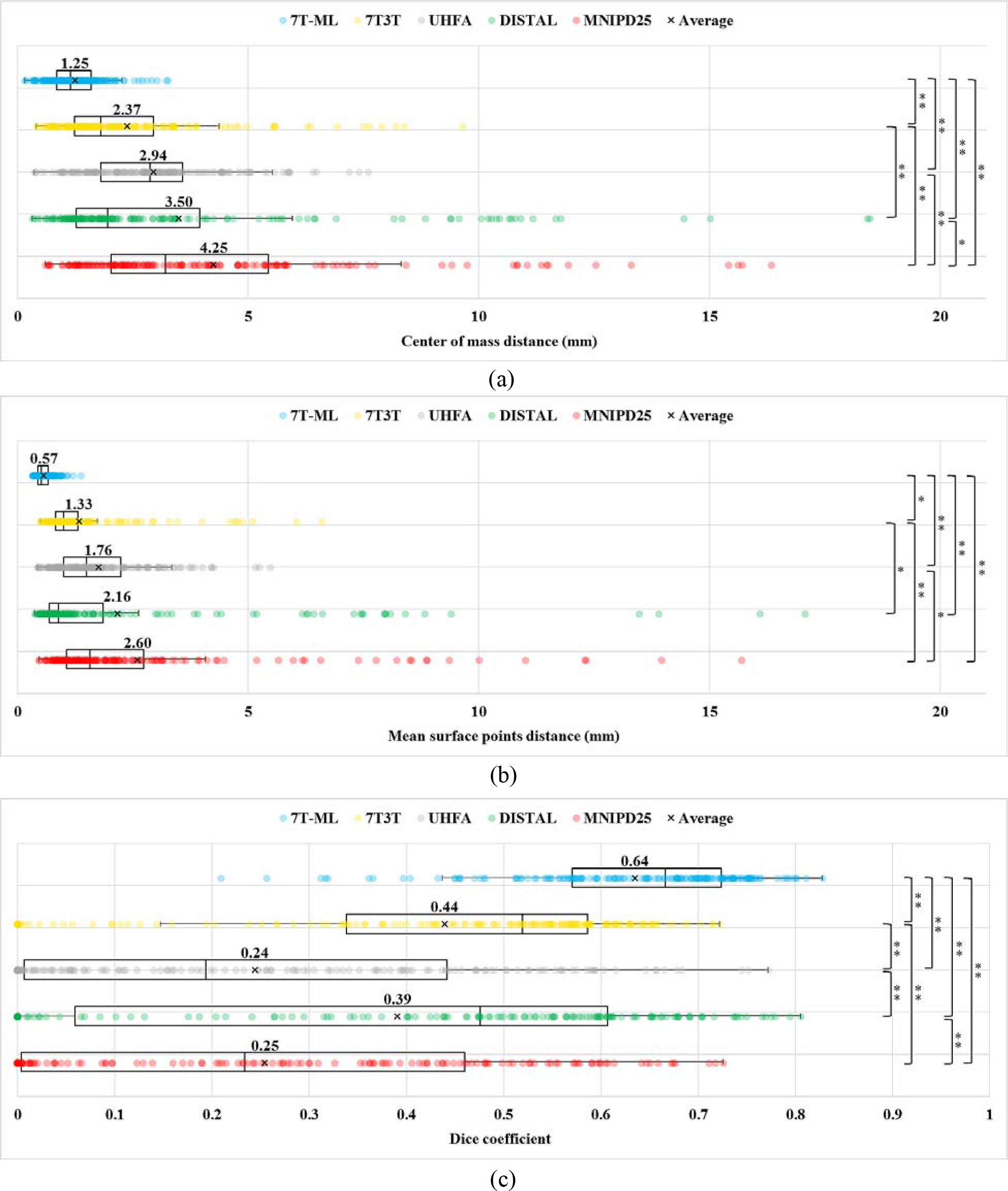
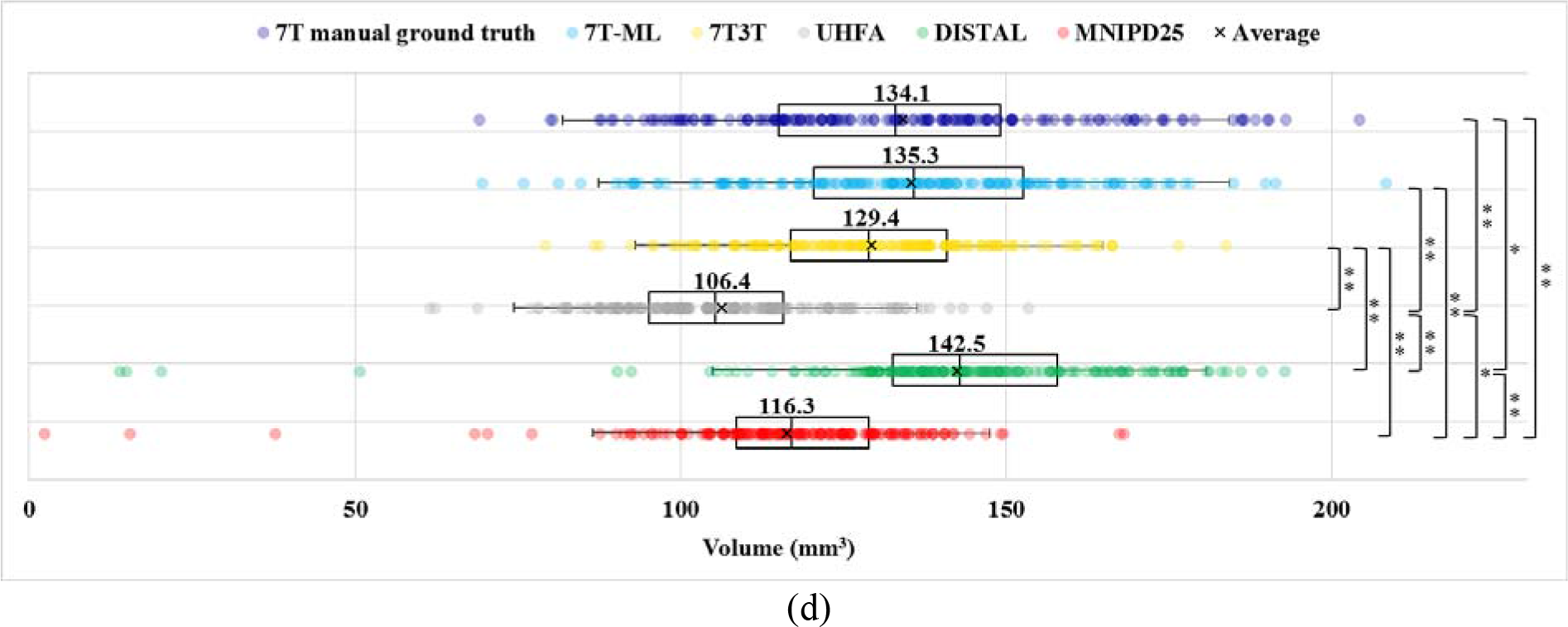
Comparison of (a) centers of mass distance, (b) mean distance of surface points (c) DC, and (d) volume for our proposed 7T-ML based STN prediction (using the bagged partial least squares regression, uniform weights, 100 ensemble size, and automatically segmented predictors), 7T3T, UHFA, DISTAL, and MNIPD25 atlases, in comparison with 7T manual ground truth across 80 patients. A one-way ANOVA and post hoc test (with Tukey’s method) is performed for multiple comparisons. The significance level is denoted by asterisks (* for p<0.05 and ** for p<0.001; ANOVA and post hoc test). Note that the volume difference between the 7T manual ground truth and our 7T-ML based STN and 7T3T, respectively, are not significant (p>0.05; ANOVA and post hoc test.)

The 7T-ML also yields significantly better average mean distance of surface points and DC values (0.57±0.18mm and 64±12%) than 7T3T (1.33±1.03mm and 44±21%), UHFA (1.76±1.01mm and 24±23%), DISTAL (2.16±2.97mm and 39±27%), and MNIPD25 (2.60±2.73mm and 26±24%) atlases, respectively, in comparison with the ground truth (p<0.0001; ANOVA).

Volumes of the 7T-ML based predicted STN (135.3±25mm^3^) and 7T3T atlas (129.4±19mm^3^) are comparable to those of the 7T manual ground truth STN (134.1±29mm^3^) on average (p>0.05; ANOVA and post hoc test), while the STNs based on other atlases are significantly different (p<0.05; ANOVA and post hoc test). DISTAL atlas-based STN is the largest (142.5±27mm^3^), but UHFA and MNIPD25 based STNs are smaller on average (106.4±23mm^3^ and 116.0±21mm^3^, respectively).

Fig. 3-(a) presents histograms of the centers of mass distance measured on 160 STNs computed by the various discussed methods. Each zone represents bins that include median or average centers of mass distance (bin size: 0.5mm). Median and average centers of mass distance of our proposed 7T-ML STN are in zone (i), while median and average centers of mass distance for the STNs computed based on standard atlases are distributed in zones (ii) and (iii). Our proposed 7T-ML demonstrates both higher accuracy and higher consistency in the centers of mass distance than standard atlases (i.e., smaller average and narrower distribution). Fig. 3-(b) and (c) visualize, for the compared methods and example subjects, the STNs in different average and median centers of mass distance zones, exemplifying the superior performance of the proposed 7T-ML.

**Figure 3:**
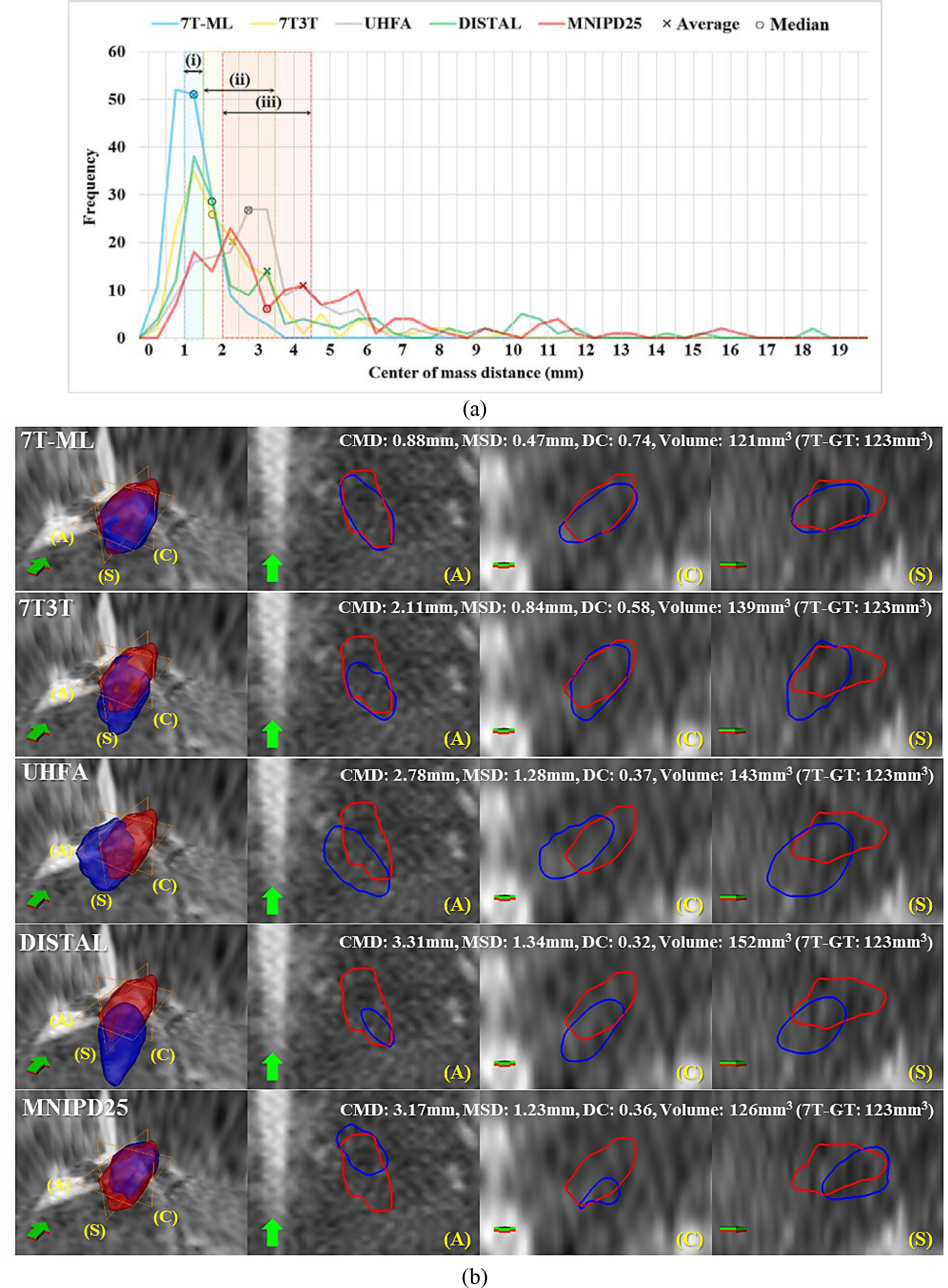
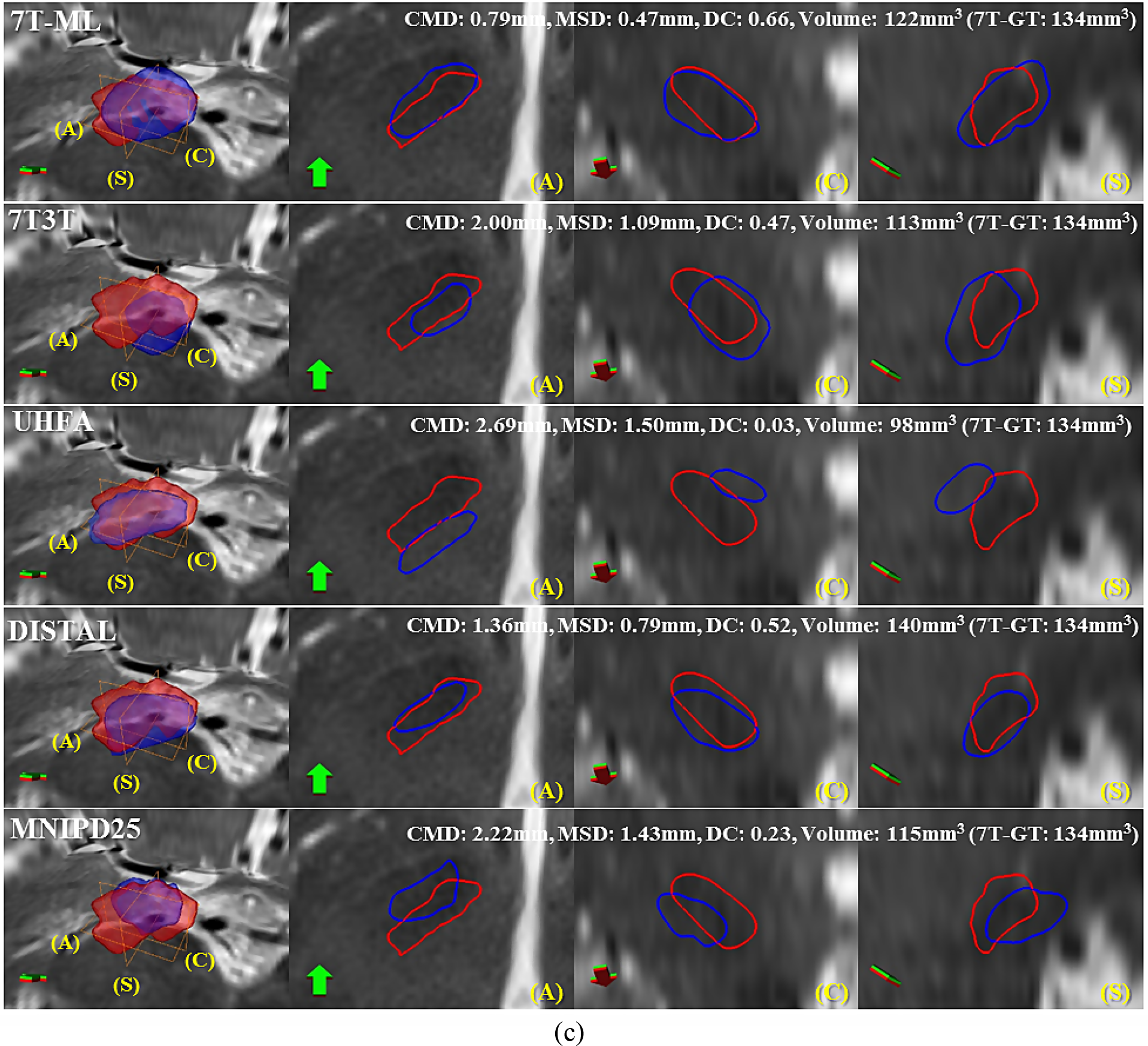
Centers of mass distances’ histogram and visual examples of the proposed 7T-ML and standard atlases (blue) overlaid with 7T manual ground truth (red) from specific subjects. (a) Comparison of histograms of centers of mass distance measured on 160 STNs obtained from the different methods. (b) Visualization of the STNs from zone (i) and (ii) in (a), representing average centers of mass distance of the 7T-ML and standard atlases, respectively on the 1.5T MRI of an example PD patient (age: 51). (c) Visualization of the STNs from zone (i) and (iii) in (a), representing median centers of mass distance on the 3T MRI of an example PD patient (age: 79). From left to right: 3D surface and contours in (A) axial, (C) coronal, and (S) sagittal planes along with arrows indicating the anterior direction. CMD: centers of mass distance. MSD: mean distance of surface points.

The STN segmentation results based on the active shape model and active appearance model framework (with 7T priors) showed 1.51±0.78mm (centers of mass distance), 0.61±0.20mm (mean distance of surface points), and 52.3±12% (DC) on average, comparing to the 7T manual ground truth. 79.4% (127/160) of the segmented STNs were less than 2mm in centers of mass distance. Our 7T-ML was still significantly better in average centers of mass distance and DC (p<0.001; ANOVA). The volume of the segmented STN was also significantly different from our 7T-ML and the 7T manual ground truth (p<0.001; ANOVA).

We also observed that the distance between active contacts and the 7T-ML STN’s centers of mass (again, here used to provide per-patient intrinsic coordinates) is much closer to that of the 7T manual ground truth STN in comparison to standard atlas-based results (Table 7; compare 2.4mm for our 7T-ML and ground truth and 3.2-5.6mm for standard atlases; see also Table S1 in the supplemental material for data on all the contacts). Once again, the STN’s center of mass is not clinically considered the optimal target position, and is here used simply as a reference for intrinsic coordinates. The target region (preferred by implanting team) can be defined/visualized within our 7T-ML STN. Nevertheless, the results here reported for our tested clinical data are comparable to those in the literature [Pallavaram et al., 2015].

**Table 7:**
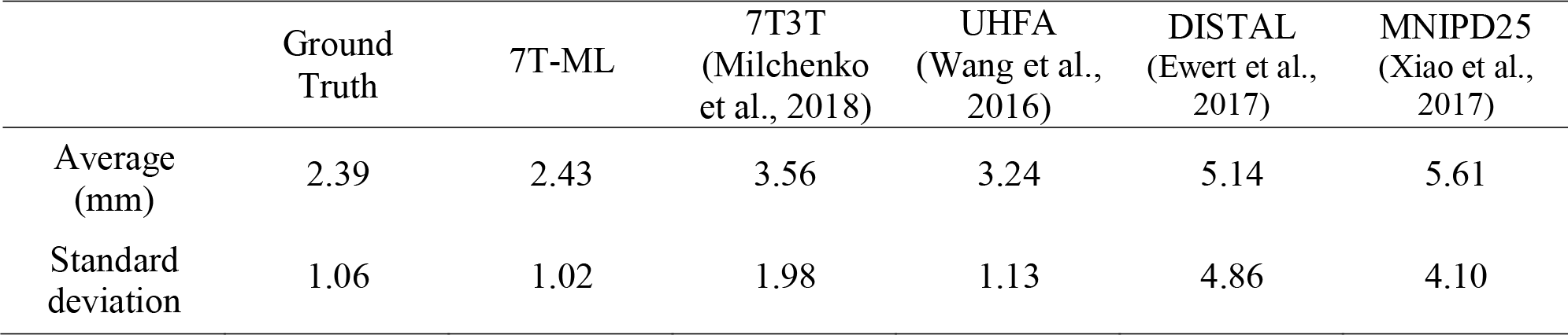
Distance between active contacts and STN’s centers of mass computed from the 7T manual ground truth, proposed 7T-ML, 7T3T, UHFA, DISTAL, and MNIPD25 atlases.

Furthermore, as shown in Table 8 and Fig. 4, the active contacts were frequently populated at the posterior lateral parts (i.e., regions 2 and 4) of the STN as obtained from the 7T manual ground truth STN. The proposed 7T-ML was consistent with this observation, while the spatial distribution of active contacts was markedly different in the atlas-based results. This is repeated for all the contacts in Table S2 and figures S1-S4.

**Figure 4:**
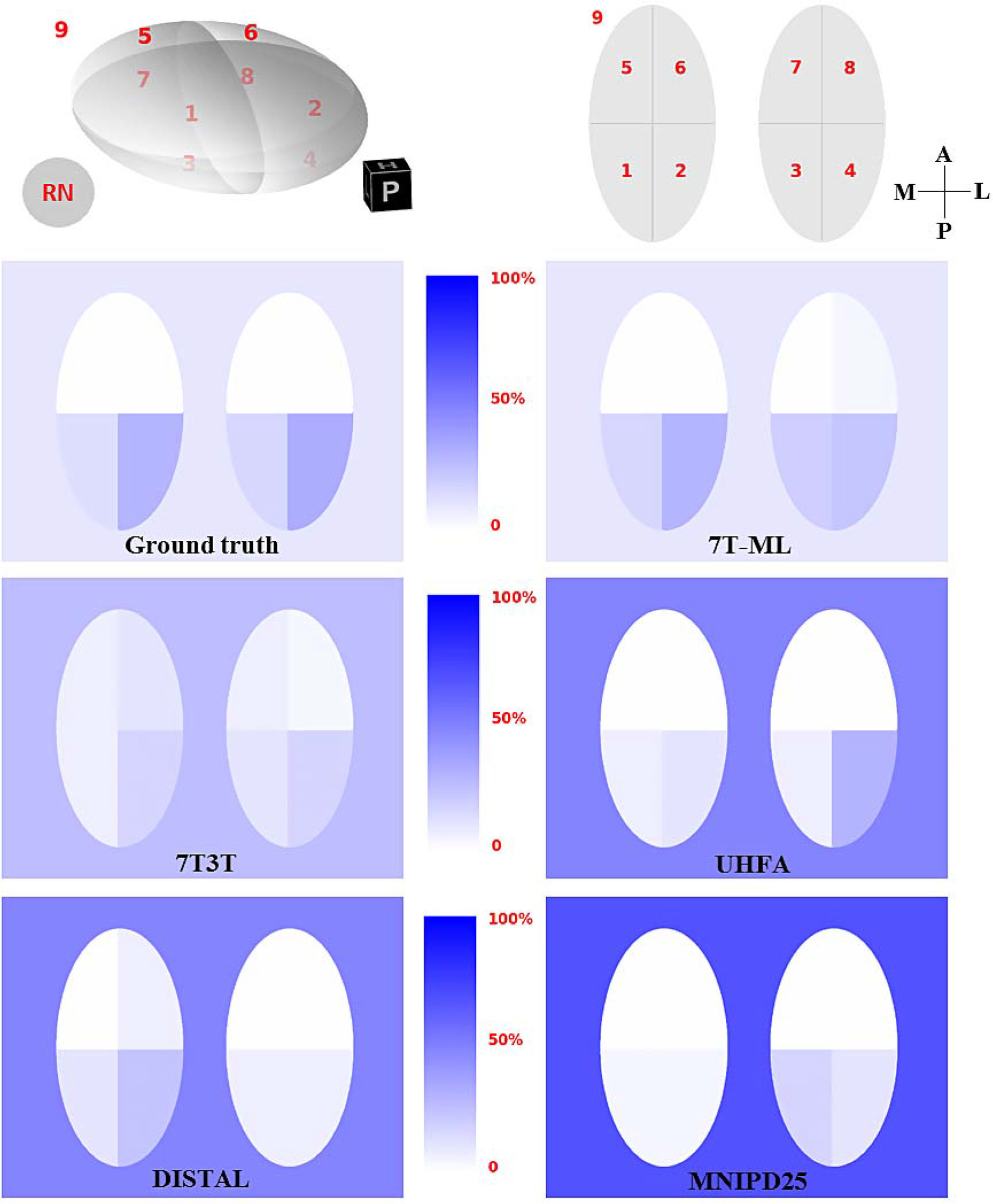
Density maps of the active contact population in different regions within the per-patient ellipsoid representation of the STN based on 7T manual ground truth, proposed 7T-ML, 7T3T, UHFA, DISTAL, and MNIPD25. Top left represents the region numbers in 3D and the right represents the corresponding numbers in the 2D slices along with posterior(P)/anterior(A) and medial(M)/lateral(L) axes. (The example shows the right STN. Left STN is mirrored into M-L direction for the population analysis.)

**Table 8:** Numbers of active contacts placed in each region within the per-patient ellipsoid representation of the STN computed based on the 7T manual ground truth, proposed 7T-ML, 7T3T, UHFA, DISTAL, and MNIPD25 atlases. See Fig. 4 for the localization of each region.

| Region | Ground truth | 7T-ML | 7T3T<br>(Milchenko et<br>al., 2018) | UHFA<br>(Wang et al.,<br>2016) | DISTAL<br>(Ewert et al.,<br>2017) | MNIPD25<br>(Xiao et al.,<br>2017) |
| --- | --- | --- | --- | --- | --- | --- |
| 1 | 4 | 5 | 2 | 2 | 3 | 1 |
| 2 | 9 | 9 | 5 | 3 | 7 | 1 |
| 3 | 5 | 6 | 3 | 2 | 2 | 5 |
| 4 | 10 | 7 | 5 | 9 | 2 | 3 |
| 5 | 0 | 0 | 2 | 0 | 0 | 0 |
| 6 | 0 | 0 | 3 | 0 | 2 | 0 |
| 7 | 0 | 0 | 2 | 0 | 0 | 0 |
| 8 | 0 | 1 | 1 | 0 | 0 | 0 |
| 9 | 3 | 3 | 8 | 15 | 15 | 21 |

Fig. 5 presents an STN visualization based on the 7T manual ground truth, our proposed 7T-ML, 7T3T, UHFA, DISTAL, and MNIPD25 atlases, along with the post-operative electrode contacts of a specific PD patient. According to the post-operative monopolar reviews, the activation of the contact 1 was associated with the best motor improvement (62%), while activation of other contacts had resulted in lower motor improvements (15-46%). Contact 1 was entirely inside the dorsal STN according to our proposed 7T-ML, confirmed with the 7T manual ground truth STN. Active contacts associated with motor improvements were also placed in the dorsal area of the STN based on 7T3T, DISTAL, and MNIPD25. However, the atlas-based STNs were associated with large errors in shape and location with respect to the ground truth (unreliable STN trajectory). Note that a system operating/deciding based on our proposed 7T-ML system would have achieved optimal results (with the predicted STN close to the ground truth and the current lead location), while the same system operating based on those atlases would have misled the electrode placement, thereby resulting in adverse effects. Therefore, atlas-based STN segmentation would require further revision before being utilized into a clinical setup. While a full investigation of these aspects and consequences is beyond the scope of this study, these results confirm that our proposed sub-region accuracy 7T-ML provides a reliable guide for the STN (sub-region) targeting for DBS surgery and treatment based on patient-specific standard clinical MR data.

**Figure 5:**
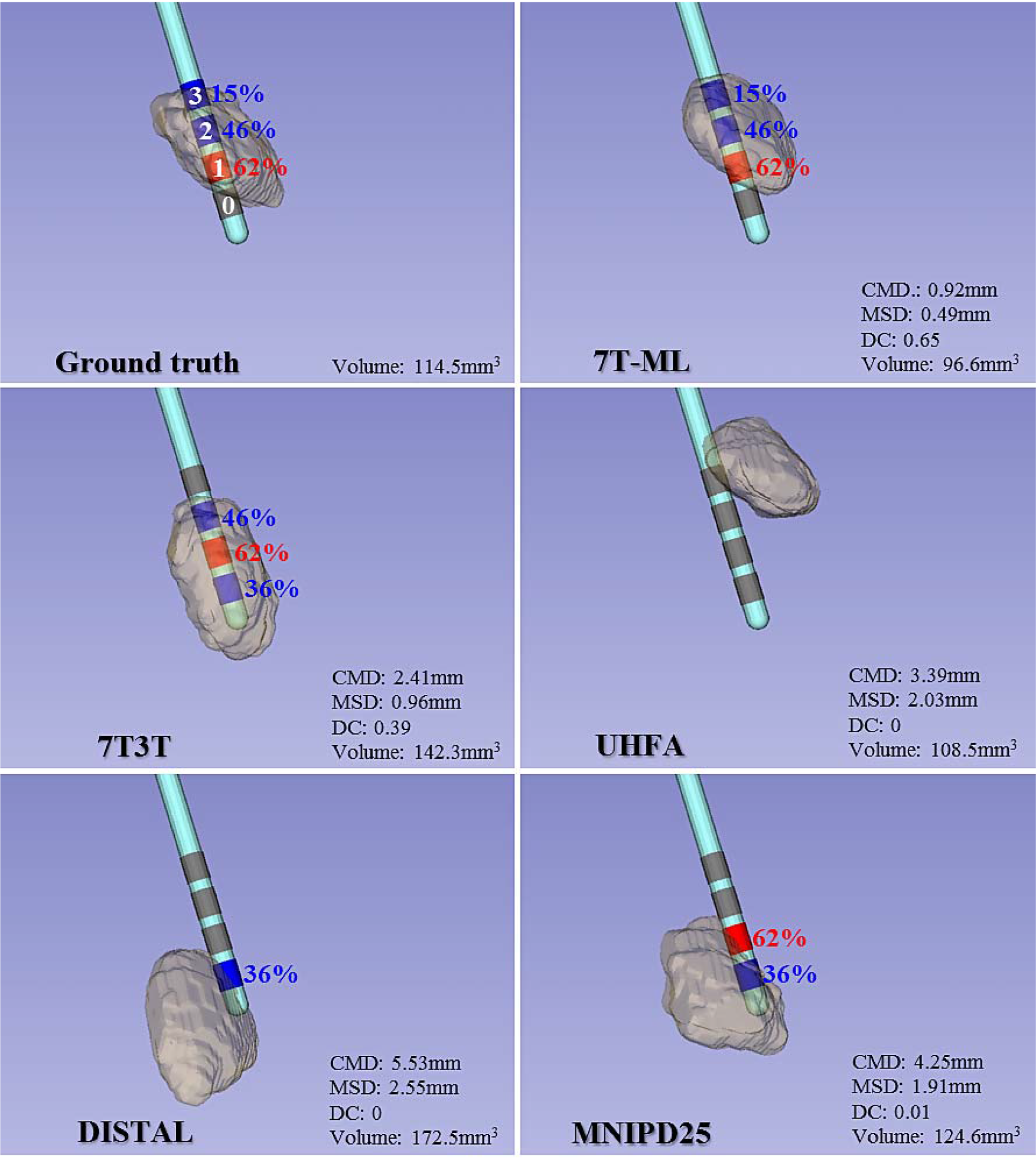
Comparison of the STN computed based on the 7T manual ground truth, our proposed 7T-ML, 7T3T, UHFA, DISTAL, and MNIPD25 atlases, overlaid with the electrode contacts for a specific PD patient (age: 48). The visualization is an example from zone (i) and (iii) in Fig. 3-(a), representing average centers of mass distance of our 7T-ML and standard atlases-based STNs, respectively. Electrode contacts were placed into the dorsal zone according to the ground truth. Similarly to the ground truth, our 7T-ML STN completely includes the active contact (red, contact 1) associated with the best motor improvement (62%) and the active contacts (blue, contact 2 and 3) with lower motor improvement (15-46%). Dorsal STNs based on 7T3T, DISTAL, and MNIPD25 also would have been activated by the contacts associated with motor improvements, but they showed much larger errors in shape and location than 7T-ML. This means that it might lead to misleading placement. None of contacts are placed in the dorsal STN based on UHFA. CMD: centers of mass distance. MSD: mean distance of surface points.

Next, the effect of the main components in our proposed 7T-ML framework is analyzed as follows:

1. *Predictors*: the average centers of mass distance between the predictors (non-STN subcortical structures) automatically segmented on the selected clinical MRI and their manual 7T MRI counterparts was 0.92±0.45mm. The 7T-ML STN using manually segmented predictors showed comparable results to these observed using automatically segmented predictors (Fig. 6; p>0.05; ANOVA).
2. *Regression methods*: The 7T-ML using the bagged partial least squares regression produced significantly better results than using the regression forest method (figures 6 and 7; p<0.05; ANOVA).
3. *Ensemble sizes*: The number of training subsets (10, 100 and 200) had insignificant effect on the STN prediction accuracy (p>0.05; ANOVA).
4. *Weighting methods*: Incorporating non-uniform weights based on estimated error scores produced comparable results to these observed using uniform weights (Fig. 7; p>0.05; ANOVA) for the bagged partial least squares regression and regression forest methods. However, incorporating the actual error scores for determining the weights of the training sets resulted in significantly better prediction with the regression forest method (p<0.01; ANOVA), but insignificant with the bagged partial least squares regression. The mean squared error computed between the estimated error scores and the actual ones was lower for the regression forest method in comparison to the bagged partial least squares regression (Fig. 8; p<0.05 ANOVA). The smaller the number of training subsets, the higher the obtained mean squared error. This observation suggests that the robust framework was more effective when using the regression forest method with over 100 subsets.

**Figure 6:**
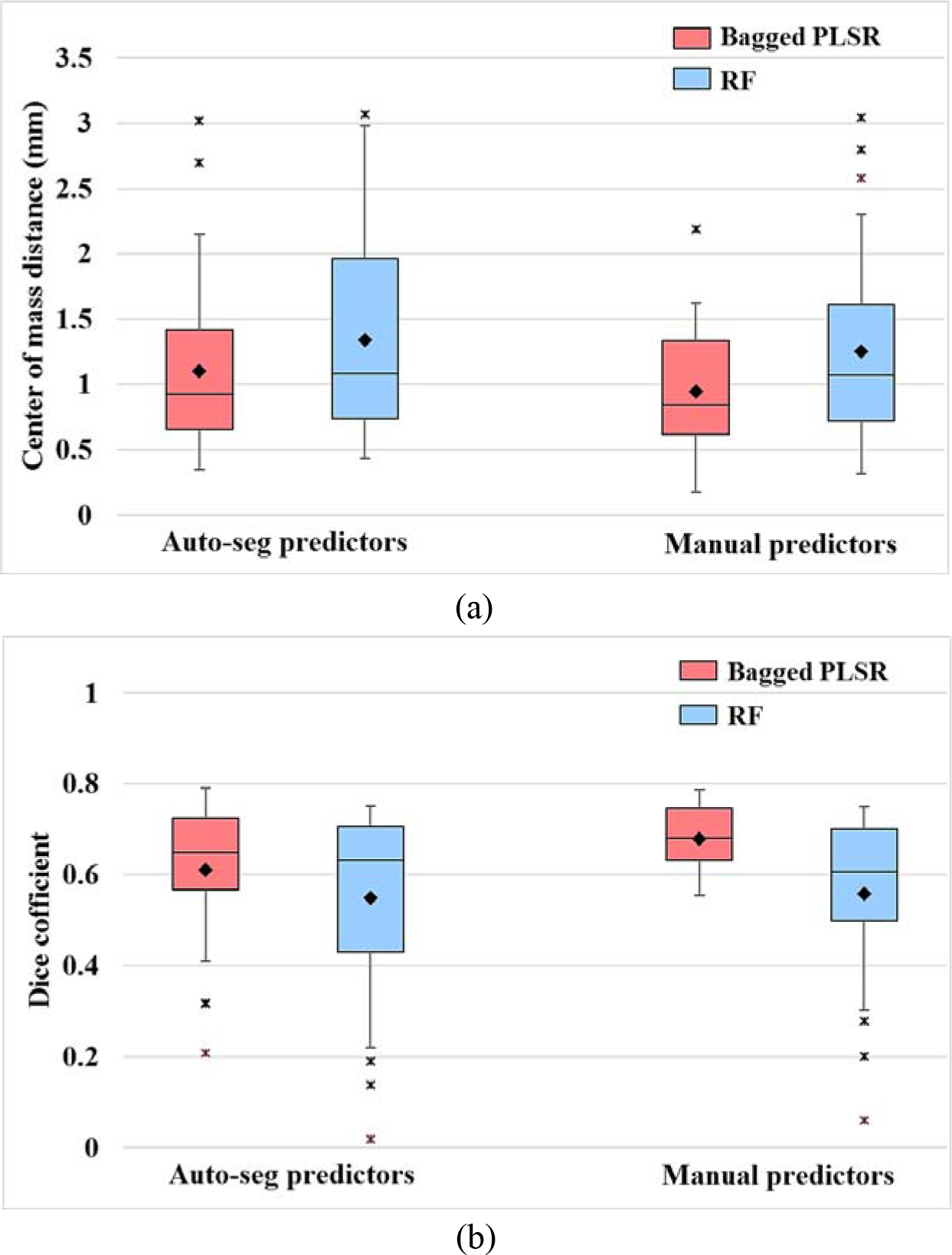
Comparison of (a) centers of mass distance and (b) DC values for predicted STNs with automatically and manually segmented predictors on the 1.5T MRI of 15 PD patients.

**Figure 7:**
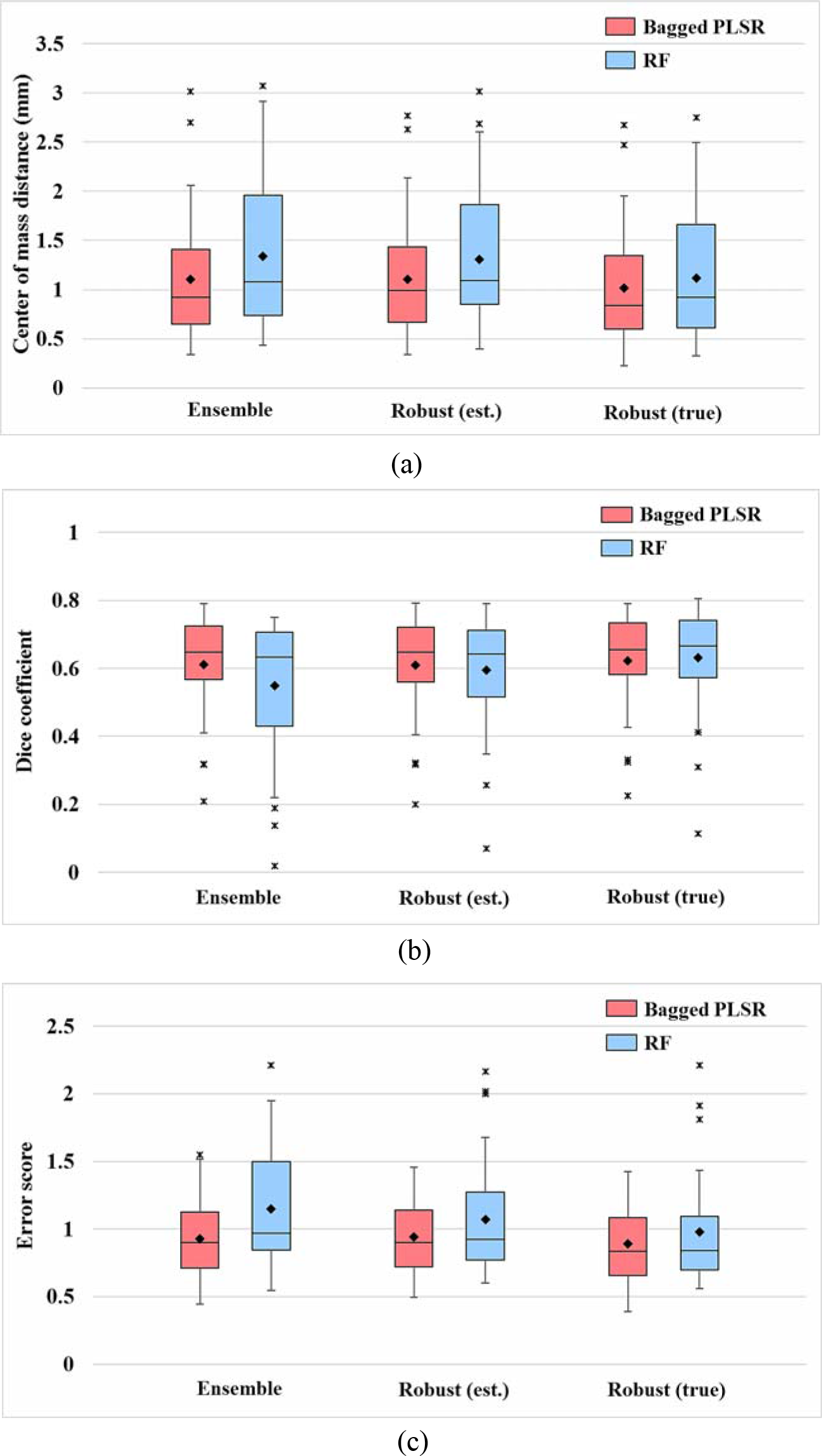
Comparison of (a) centers of mass distance, (b) DC values, and (c) error scores for ensemble STN prediction and robust prediction (for estimated error scores and actual ones) by bagged partial least squares regression and regression forest learning, with automatically segmented predictors and an ensemble size of 100.

**Figure 8:**
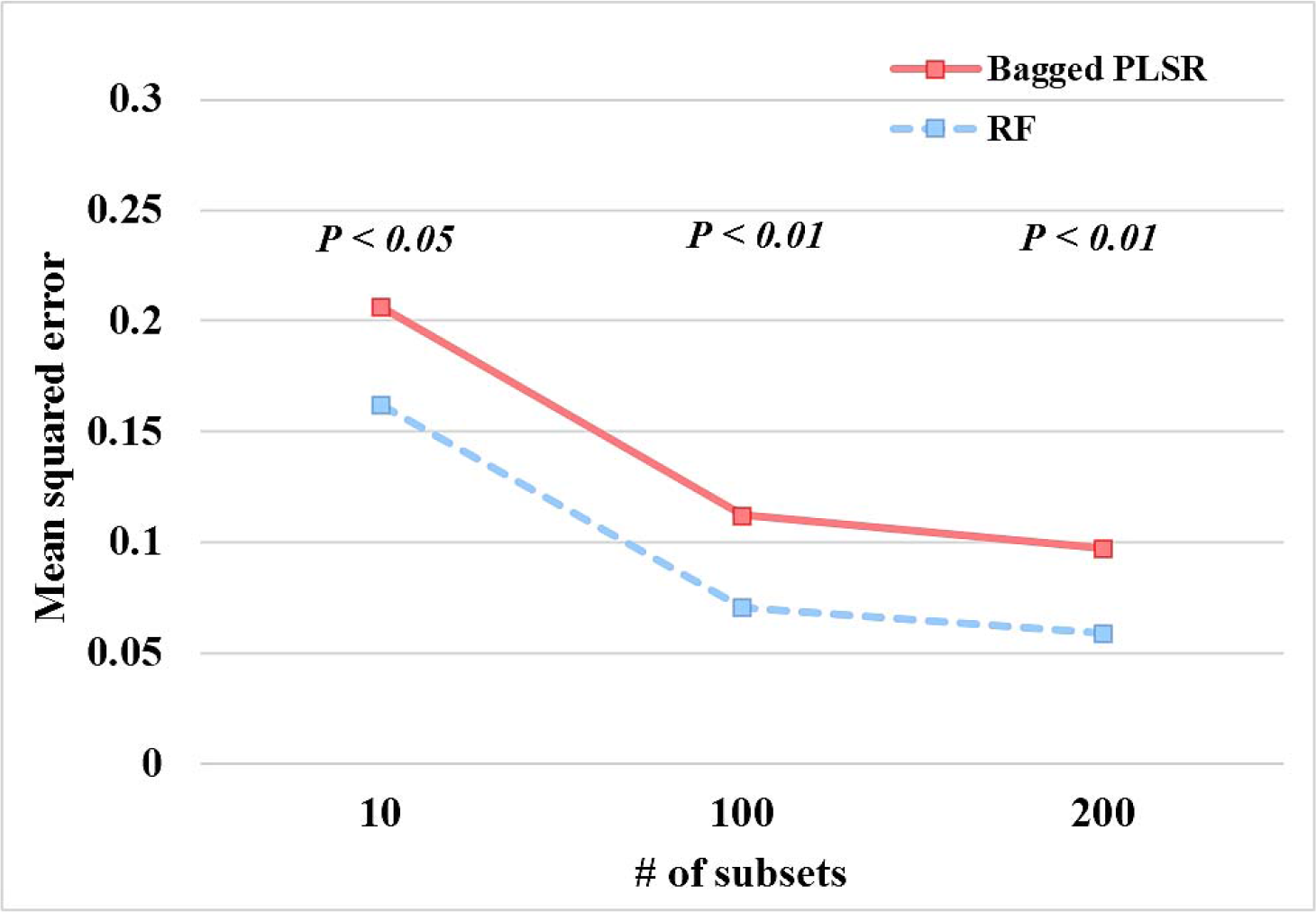
Average mean squared error between estimated error scores and true ones (computed from the 7T manual ground truth STN) for predicted STN using the bagged partial least squares regression and the regression forest according to different number of training subsets within the robust framework.

## 4. Discussion

In this work, we proposed a computational framework to automatically localize and visualize the STN based on the standard clinical MRI, where it is not clearly identifiable, by taking advantage of our 7T MRI database and machine learning. For validation, we used the STN ground truth that was defined on the 7T MRI, with careful manual annotation and cross-validation. Particularly, the 7T manual segmentation was shown to be accurate and consistent with neurophysiological data [Duchin et al., 2018; Shamir et al., 2018]. Moreover, the most-effective contacts were located in the dorso-lateral area of the 7T manual STN segmentation, which matches other clinical reports [Garcia-Garcia et al., 2016; Herzog et al., 2004]. These studies show that the 7T manual segmentation data is highly consistent with biological and clinical measures. Once segmented accurately, the 7T manually segmented STN can be transformed to a lower-resolution clinical image as done here to train the proposed 7T-ML method, and still accurately represent the STN, although it may be hard to observe it on the standard clinical image. The proposed 7T-ML based STN is highly consistent with the 7T manual ground truth in center of mass, mean surface points, DC, and volume. Moreover, the population and location of the active contacts at the different subregions within the proposed 7T-ML based predicted STN highly agrees with that of the 7T manual ground truth. We believe that the consistency of our proposed method with neurological measures and clinical outcomes supports its use to facilitate the guidance of the DBS electrode based on direct targeting using standard clinical MRI of individual patients.

As summarized in Table 9, a variety of automatic STN segmentation methods based on MRI have been proposed. While most methods are not publicly available, state-of-the-art atlases and corresponding templates were freely downloadable, and thus we have tested them on our own clinical datasets. While some automatic approaches exploit the sufficient intensity information from contrast enhanced MR sequence or even higher field MRI (e.g., QSM, FGATIR, or 7T), it remains to be investigated if these methods provide accurate segmentation of the STN on standard clinical T_2_W MRI.

**Table 9:** Summary of automatic STN segmentation methods based on MRI.

| Author (year) | Method | Test data (subject-specific) |  |  |  | Implementation availability |
| --- | --- | --- | --- | --- | --- | --- |
|  |  | MRI Modality | Magnetic field strength | Dataset source | The number of datasets (clinical details) |  |
| Ewert et al. (2017) | single atlas propagation | T<sub>2</sub> | 1.5T/3T | IXI Dataset | 22 (normal) | 3T T<sub>2</sub> template (MNI)/atlas |
| Wang et al. (2016) | single atlas (7T) propagation | T<sub>2</sub> | 1.5T | Own dataset | unknown (normal) | 7T T<sub>2</sub> template/atlas |
| Xiao et al. (2017) | single atlas propagation | - | - | - | - | 3T T<sub>2</sub>* template (MNI)/atlas |
| Milchenko et al. (2018) | single atlas (7T) propagation | T<sub>2</sub> | 3T | Own dataset | 56 (PD) | 3T T<sub>2</sub> template (MNI)/atlas |
| Haegelen et al. (2012) | multiple atlases propagation (patch based label fusion) | T<sub>1</sub>/T<sub>2</sub> | 3T | Own dataset | 10 (PD) | No |
| Xiao et al., (2014a) | multiple atlases propagation (patch based label fusion) | T<sub>1</sub>-T<sub>2</sub>* fusion | 3T | Own dataset | 10 (PD) | No |
| Xiao et al., (2014b) | multiple atlases propagation (majority voting) | T<sub>1</sub>/T<sub>2</sub> | 3T | Own dataset | 10 (PD) | No |
| Plassard et al. (2017) | multiple atlases (7T) propagation | T<sub>1</sub>/Inversion recovery scan (O-IR) | 3T | Own dataset | 9 (Normal) | No |
| Bernard et al., (2012) | active shape model | SWI | 3T | Own dataset | 24 (unknown) | No |
| Kim et al., (2014) | active contour with multi-contrast edge maps and shape priors and non-overlapping constraints | SWI/T<sub>2</sub> | 7T | Own dataset | 6 (Normal) | Available upon request |
| Li et al. (2016) | level set (chan and vese with MLE initial contour) | T<sub>2</sub> | 3T | Own dataset | 10 (PD) | No |
| Garzon et al. (2017) | spatial priors based GMM of image intensities | QSM | 3T | Own dataset | 40 (Normal) | Yes |
| Visser et al. (2016b) | MRF shape prior model and intensity model | T<sub>2</sub>*(QSM) | 7T | Own dataset | 53 (Normal) | No (will be available in upcoming version of FSL) |
| Milletari et al. (2017) | Hough-voting to acquire mapping from CNN features | QSM | unknown | Own dataset | 55 (unknown) | No |
| Proposed 7T-ML | Statistical shape and pose relationship learning from 7T priors | T<sub>2</sub> | 1.5T/3T | Own dataset | 80 (PD/ET/Normal) | Available soon |

For this reason, we compared the performance of the proposed approach with that of state-of-the-art atlas-based methods: 1) 7T3T [Milchenko et al., 2018], 2) UHFA [Wang et al., 2016], 3) DISTAL [Ewert et al., 2017; Horn et al., 2017b], and 4) MNIPD25 [Xiao et al., 2017]. While these atlases reasonably localize and visualize the STN on subject-specific data they represent well, their accuracy and consistency on our tested clinical data were much lower than the proposed 7T-ML method. A one-way ANOVA and post hoc test with Tukey’s method showed that our 7T-ML is significantly better than any other popular method we tested against. Moreover, the atlas-based STNs missed a large portion of postoperative active contacts, potentially resulting in sub-optimal planning, programming, or outcomes.

Standard atlases that are well defined in a normalized space could be of great value for retrospective population studies in the field [Horn et al., 2017a; Horn et al., 2017b; Horn et al., 2017c; Keuken et al., 2014]. However, uncertainty in registration needs to be addressed when using such atlases for patient-specific STN-DBS targeting. Single atlas-based methods heavily rely on the registration quality between the atlas template and the clinical MRI from individual patients. Since the registration error may be larger on standard clinical images where the STN is not clearly visible or when the atlas template does not represent subject-specific data, atlas-based methods usually require further revision of the segmentation. All the significant errors from standard atlases are not simple biases, which would be easy to correct, the errors are unpredictable and have large variance as well. Large variability of atlas-based results on our tested clinical data might explain this issue. Furthermore, an inaccurate definition of the STN in the atlas template may produce an additional error in patient-specific targeting [Ewert et al., 2017].

We provided comprehensive results from a variety of state-of-the-art atlases to discuss uncertainty in registration that might be induced by (1) morphological variability (normal vs. patient), (2) different contrast (magnetic field or modality) between atlas template and clinical data, and (3) its suboptimization. This also confirms the benefits of our proposed 7T-ML that combines accurate STN models and machine learning for prediction from clinical data.

More specifically, the DISTAL atlas [Ewert et al., 2017] and UHFA [Wang et al., 2016] were defined on the atlas template from normal subjects, and segmentation results on data from normal subjects were provided. The results on the PD-specific data might be deteriorated by morphological variability between the atlas template and the clinical data. Moreover, UHFA [Wang et al., 2016] utilized the 7T T_2_W MRI atlas template, and thus the contrast discrepancy between the 7T MRI and standard clinical data also might have affected the registration (this might also explain the often smaller STN volume found when computed with this atlas compared with other atlases-based results).

Although the MNIPD25 atlas [Xiao et al., 2017] is PD-specific, the MR modality of the STN atlas templates (T_2_*W) is different from that of our standard clinical T_2_W data. This might have caused an error in registration between the atlas template and the clinical data; this atlas was associated with worse performance and smaller volume than using other atlases.

Milchenko et al. [2018] created a 7T atlas from elderly subjects and registered it onto the average template representing 3T T_2_W MRIs from PD patients. While promising results on PD patient-specific data was reported, segmentation results on our own clinical data (mostly PD patients) were insufficient for clinical utilization (although it showed better performance than using other atlases). Sub-optimization in a single warp might have affected the results.

Table 10 presents quantitative comparison for centers of mass distance and DC of the STNs obtained using each method, clustered according to magnetic field strength. Specifically, 7T3T and DISTAL using a 3T T_2_W MRI template, and UHFA using a 7T T_2_W MRI template, showed much better performance on 3T MRIs, that have closer appearance to those templates, than on 1.5T MRIs. This illustrates that atlas-based methods require a template image closer to a given clinical MRI to improve their accuracy. Note that our 7T-ML shows comparable accuracy regardless of the quality (and strength) of the clinical MRIs, although the algorithm was not trained on 3T MRIs. In a complementary study, we have computed with the exact same method here introduced the STN on 3T MRIs obtained in another center and observed that it is very similar (~1mm) to the STN that was defined based on MER and blindly compared to our method [Shamir et al., 2018]. These complementary studies show the high accuracy and consistency of the proposed computational method, regardless of the proxy used for representing the ground truth (7T or MER).

**Table 10:**
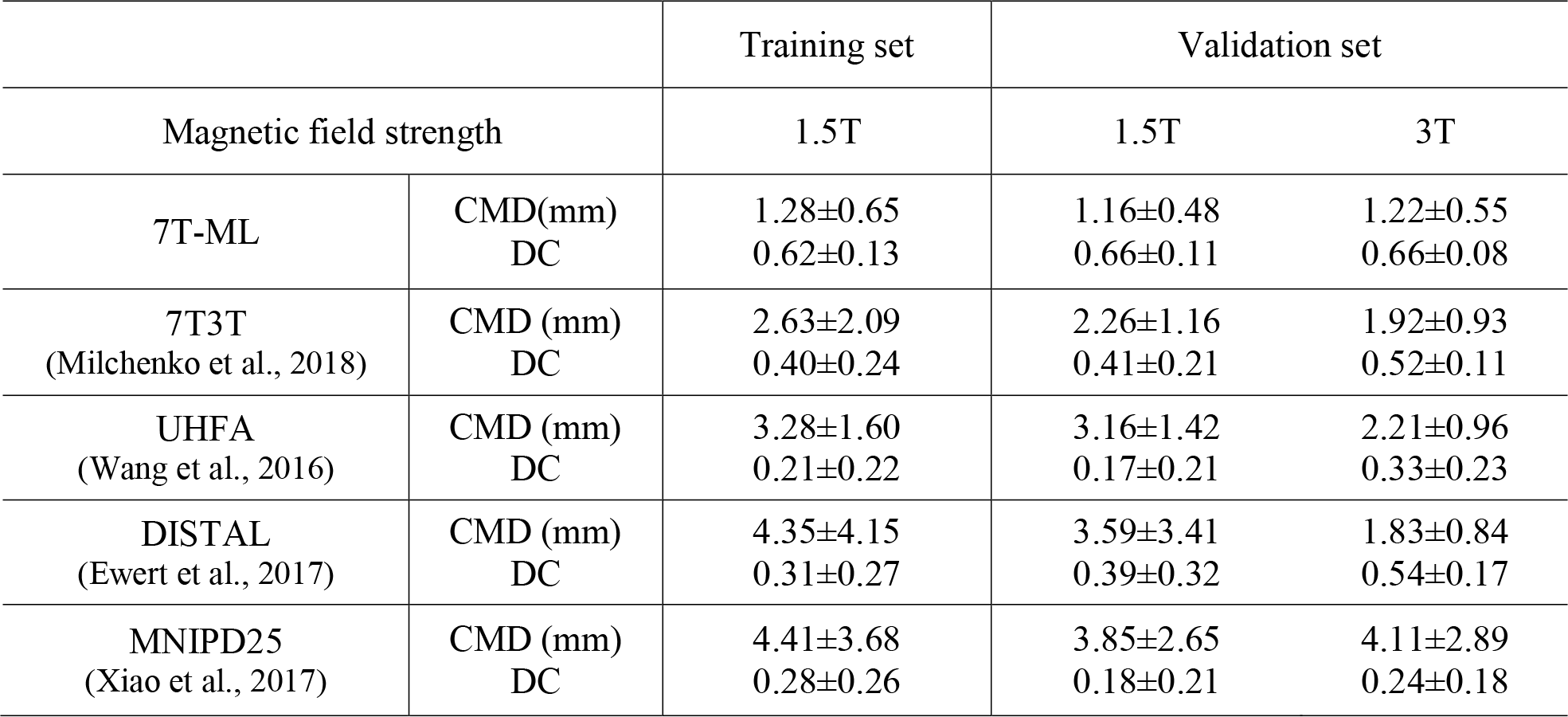
Centers of mass distance and DC value between the STNs obtained using the proposed 7T-ML, 7T3T, UHFA, DISTAL, MNIPD25 atlases and the 7T manual ground truth STNs, grouped according to magnetic field strength in the training and validation sets. A one-way ANOVA and post hoc test (with Tukey’s method) showed that the proposed 7T-ML is significantly better than atlas-based methods in each measure (p<0.0001).

Registration steps used in this work oftentimes caused large errors in atlas-based results that could lead to misplacement of the stimulating electrode and ineffective DBS treatment. An optimization of the registration processes can considerably improve the fitting of the atlas and the STN segmentation accuracy [Pallavaram et al., 2015]. However, assuring such an optimization in the single registration mode still remains a challenging task even though the field has progressed [Viergever et al., 2016]. Moreover, it is hard to generalize an optimized registration into cases in large-scale population. The full investigation of the registration performance is beyond the scope of this work, but is important for automatic targeting as here illustrated.

Our 7T-ML minimizes the effort to find the optimal single warp in a robust and fully automatic way. More specifically, we register clinical training images in our database directly to the clinical query image and select the most similar sets to reduce random error and bias in the single warp [Aljabar et al., 2009]. Moreover, we refine the STN location and shape by learning 7T knowledge. It should be noted that although our 7T-ML framework uses the same registration steps that affected inaccurate atlas-based segmentation, thanks to these important additions the obtained results are significantly better than those obtained using the standard atlases.

Recent multiple atlases-based approaches showed promising results for the localization of the STN [Haegelen et al., 2013; Xiao et al., 2014a; Xiao et al., 2014b]. In these studies, manual labeling was performed, based on the appearance of 3T MRI, and automatic segmentation of the subcortical structures on the query patient was done by registration between the 3T MRIs atlas templates and patient. While the automatic segmentation closely matches its manual STN on the patient images, taking advantage of multiple atlas templates with similar appearances (reducing the registration uncertainty), it is unclear if the STN that appeared as hypo-intense on the 3T MRI reflects an accurate geometrical representation of the STN of individual subjects, especially in the dorso-lateral part that is critical for DBS targeting [Cho et al., 2011; Plantinga et al., 2014] (see also difference between volumes of 7T manual ground truth and MNIPD25 atlas-based STN in Fig. 2-(d)). Moreover, employing such methods in standard clinical scenarios based on lower quality images (e.g., 1.5T MRI) may result in larger errors since the registration accuracy is expected to be lessened [Avants et al., 2011b; Ou et al., 2014].

Using current standard clinical imaging protocols, it is not feasible to differentiate the STN from the SN in clinical 1.5T or 3T MRI [Abosch et al., 2010]. As such, it remains unclear how approaches based on intensity and texture information on the target image [Bernard et al., 2012; Li et al., 2016] handle leakage around the border between the STN and SN. To explore this issue, we automatically segmented the STN using the active shape model and active appearance model framework (the same method that was used for segmentation of predictor structures in our 7T-ML framework). While it reasonably localized the STN with the initialization from the 7T priors, our 7T-ML was still significantly better in average centers of mass distance and DC. Particularly, the size of the segmented STN was much smaller than the 7T manual ground truth. This might be attributed to fuzzy boundary of the STN on the image.

Some approaches use pre-processed sequences or high quality data to make the STN more discernible or segment visible various mid-brain structures. For example, Garzón et al. [2017], Visser et al. [2016b], Milletari et al. [2017], and Plassard et al. [2017] utilized contrast enhanced MR sequences (QSM or FGATIR) to visualize subcortical structures, including the STN. Garzón et al. [2017] mentioned that the algorithm showed lower accuracy on R_2_* and T_2_W FLAIR images, indicating that it was specialized to high contrast data. Visser et al. [2016b] automatically segmented the STN on the 7T multimodal MRI. Also, Visser et al. [2016a] segmented the striatum and globus pallidus - that are fairly visible on 1.5T T_1_W MRI. Recently, several state-of-the-art methods using deep neural networks to segment brain structures are of interest [Bao and Chung, 2015; De Brébisson and Montana, 2015; Dolz et al., 2017; Shakeri et al., 2016]. Similarly, they focused on segmentation of brain regions (e.g., Thalamus, Caudate, Putamen, Pallidum, etc.) that are discernable on the image. Low quality clinical images, where even manual segmentation of the STN is not possible, might lead to challenges when using deep learning for the task here considered [Zhou et al., 2018]. We could potentially apply deep learning architectures to instead segment the predictor structures (fairly visible on the clinical image) that are used to predict the STN.

The proposed 7T-ML leverages our 7T MRI database and machine learning to predict the STN that is not normally visible on the clinical MRI. It learns anatomical knowledge encoded from our 7T training data, which are independent of image intensity values. The 7T-ML, thereby, achieves comparable results to 7T manual segmentation on the clinical image. We observed an average of 1.1±0.6mm in centers of mass distance (93% of the cases were better than 2mm accuracy) from even lower quality data (1.5T MRI) of selected 15 PD patient’s data used for the in-depth study. This is consistent with the reported results for 160 STNs from 80 subjects and also validates that the 7T-ML is robust on clinical 1.5T MRI as well as 3T MRI.

Similarly to the 7T manual ground truth, the 7T-ML based STN accurately localized and predicted the active contacts, especially in the posterior lateral region that is considered a motor territory often targeted for Parkinson’s DBS [Plantinga et al., 2016]. We should stress that we analyzed the spatial relationship between contact location and the STN using the ellipsoid representation within the STN in an individual patient data space. This is important since the transformation of the patient data into the common space and vice versa entails biases in the relationship. This also validates that our proposed 7T-ML approach facilitates precise localization of the electrode’s leads within patient-specific STN’s subregions. A few active contacts were found to be outside the posterior part of our 7T-ML and the 7T manual ground truth STN. However, all of them were placed closely to the boundary, possibly having an overlap between the volume of tissue activated and the STN sub-region. Small resampling or registration errors may also explain this slight mismatch. Micro-lesion effect may bias contact localization as well [Granziera et al., 2008]. Overall, our results show that while standard atlases do not achieve the accuracy and consistency needed for sub-region STN targeting, the proposed 7T-ML is successful in this major DBS challenge: consistently providing accurate target localization. Importantly, the clinical feasibility of our 7T-ML approach is further demonstrated, comparing to MER mapping in Shamir et al. [2018].

The 7T-ML STN resulted in what could be considered at first glance as relatively low DC values, although the geometric measurements are within the tolerance level of the stereotactic frame used for the surgery [Shamir et al., 2009]. The partial volume effect for small structures, such as the STN, under clinical imaging resolution affects the DC values [Hoffman et al., 1979]. It has been also reported that the size of objects affects the DC, where small structures are associated with smaller DC values [Rohlfing, 2012; Zou et al., 2004]. Shamir et al. [2016] provided numerical analysis of DC by modeling the clinical MRI resolution and center of mass error distribution in the STN manually segmented on the 7T MRI, and an average upper bound DC was estimated at 64%. This indicates that the proposed 7T-ML, with average 63.5% DC value, achieves near optimal accuracy. Recent automatic segmentation of the STN on the 7T data also resulted in comparable DC values [Visser et al., 2016b]. We should note that our 7T-ML targeted the STN on only standard clinical T_2_W MRI data where its borders are not visible, while the automatic segmentations in the above studies were obtained from the 7T multi-modal MRI with clear texture and boundary information.

The geometric distortion on the 7T MRI and inaccurate co-registration between the 7T MRI and clinical data may potentially affect the quality of the 7T priors and thus cause errors in our 7T-ML framework. Therefore, minimizing such biases was critical to increase the reliability. Experts in the team manually segmented the STN and it predictor subcortical structures by leveraging superior contrast and anatomical details on both 7T T_2_W and SWI and carefully cross-validated. Furthermore, Duchin et al. [2012] demonstrated clinical feasibility of the 7T MRI by evaluating the distortion based on the coregistration quality between 7T and 1.5T MRI. Following the proposed protocols we performed the coregistration between 7T and clinical MRI. Recently, we further validated that the 7T manual STN segmentation is highly consistent with the MER data [Duchin et al., 2018; Shamir et al., 2018].

We also examined the effect of multiple factors in our proposed 7T-ML approach. Generally, and as expected, the accuracy of predictors’ segmentation highly affects the accuracy of the resulting STN prediction. We observed comparable STN prediction results using the manual and automatic predictors’ (non-STN) segmentation. This indicates that an error level in predictors’ segmentation on the clinical data was not influential in the STN prediction, and the automatic segmentation was near optimal. Comparable STN prediction accuracy was also observed for 10, 100 and 200 randomly selected subsets from the training set, but the estimation of STN prediction error was more accurate as the subset size increased. Weighting the training set based on the estimation of its contribution to the prediction accuracy reduced the variance observed with regression forest, but not bagged partial least squares regression. Therefore, the larger subset size results in more accurate estimation of STN prediction error that, in turn, helps to improve the STN prediction (an ensemble size of 100 is considered sufficient and used for validation of our 7T-ML approach). Adding more subjects to our database may result in a significant correlation between the subset size and the STN prediction accuracy, which was not yet observed with training sets.

## 5. Future Work and Conclusions

We are currently investigating the shape refinement of the 7T-ML STN on a standard clinical MRI. If the ventral border of the STN that is adjacent to the SN can be identified in an automatic way, the STN prediction can be further refined to facilitate an even more reliable targeting. However, it remains questionable if the intensities around the STN boundary are consistent across clinical MR datasets from a large population of patients and centers.

While we focused on localization of the STN in this manuscript, which is the most popular DBS target for Parkinson’s disease, the approach presented here can be exploited for segmenting other structures such as the internal globus pallidus and Vim. With a greater number of centers beginning to target the internal globus pallidus for PD and given that it is the predominant target for dystonia, precise localization of the internal globus pallidus and its sub-regions may also prove valuable for physicians targeting this structure.

The identification of the STN on a standard clinical MRI is challenging. Therefore, more than one targeting method is incorporated today in DBS practice, often involving a more time-consuming and potentially extended-risk approach using intraoperative validation of electrode location with microelectrode recordings. Given that MER requires a level of expertise not typically found in most surgical centers, its utility and the ability of surgical sites to localize the STN and its sub-regions using this technique is highly variable across centers. To address these problems we introduced a patient-specific automatic software-only method for the visualization of the 3D STN location and shape from standard clinical MRI. The method incorporates a database of high-field 7T MRI and a novel set of machine learning algorithms. The experimental results validated that our proposed 7T-ML approach can automatically and accurately localize the STN and its sub-regions on standard clinical MRI. This work provides neurosurgeons and neurologists with accurate means for automatic patient-specific targeting of the STN and its sub-regions, potentially reducing the need for other approaches that may lengthen the procedure and/or be associated with a higher risk of side effects. Surgical Information Sciences, Inc. plans to make the 7T-ML based STN segmentation tool available for its clinical use in the near future.

## Acknowledgements

This study was partially supported by the NIH R01-NS085188, P41 EB015894, P30 NS076408, the University of Minnesota Udall center P50NS098573, NSF, ARO, ONR, and NGA.

